# Emergence of Balanced Cortical Activity via Calcium-Regulated Synaptic Homeostasis

**DOI:** 10.1101/2025.09.04.674182

**Authors:** Carl van Vreewsijk, Farzada Farkhooi

## Abstract

Cortical circuits must stabilize activity while retaining the variability and flexibility essential for computation. This raises a fundamental question: how can excitatory (E) and inhibitory (I) synapses co-adapt through homeostatic plasticity without disrupting network function or relying on fine-tuned parameters? We propose a solution grounded in the multidimensional nature of intracellular calcium signaling, which independently regulates protein synthesis at E and I synapses. By analytically characterizing calcium dynamics driven by spike-train statistics, we show that calcium’s mean encodes firing rate while its variance reflects spike-time irregularity—two complementary features critical for stable yet flexible spiking. Leveraging this dual signal, we construct a closed-loop model in which inhibitory synapses are regulated by calcium’s mean and excitatory synapses by its variance through independent pathways. This mechanism preserves irregular spiking and stabilizes firing rates across diverse inputs. Strikingly, it also yields the empirically observed weakening of synaptic strengths with the number of inputs *K* as 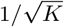, leading to the spontaneous emergence of balanced excitatory–inhibitory dynamics. These results uncover a calcium-driven regulatory principle linking intracellular signaling to the origin of balanced activity in cortical networks.

## 1 Introduction

Cortical circuits operate in inherently dynamic environments, continually shaped by experience-dependent plasticity, developmental refinement, and ongoing molecular turnover. Yet despite this flux, they maintain remarkably stable functional dynamics, with conserved patterns of spiking activity observed across species and physiological states [MOS^+^16]. A key factor underlying this stability is the homeostatic co-regulation of excitation and inhibition (E/I): neurons must dynamically adjust both excitatory and inhibitory inputs to stabilize their firing rates and prevent pathological states such as epilepsy [Tur04, HBT^+^05]. This process is thought to be governed by homeostatic plasticity mechanisms that sense neural activity and modify synaptic strengths via feedback regulation [TLD^+^98].

However, a fundamental theoretical challenge remains unresolved: how can neurons stably co-regulate both E and I synapses using feedback from a shared postsynaptic activity signal? Classical homeostatic models typically rely on negative feedback to stabilize a neuron’s average firing rate [AL93]. While these mechanisms are successful when only one synapse type is plastic, they tend to destabilize when both E and I synapses are allowed to adapt simultaneously. As we show in Section 2.1, such dual co-regulation leads to an overdetermined system in which the synaptic updates compete rather than coordinate, causing runaway dynamics unless their set points are precisely matched—a condition unlikely to be met in biological systems.

To avoid this instability, previous approaches have relied on various constraints, such as holding one synapse type fixed [AL93, VSZ^+^11], enforcing mismatched timescales for E and I plasticity for finite-time stability [DA05], or introducing external balancing forces [LMA93]. Yet these solutions limit the autonomy of local homeostasis rules and conflict with experimental evidence that E and I synapses change in parallel, regulated by distinct intracellular pathways [SM06, PZS^+^10].

To resolve this challenge, we propose a biologically grounded mechanism based on the multidimensional nature of intracellular calcium dynamics. Calcium is a canonical integrator of postsynaptic activity, sensitive to both the rate and pattern of spiking [IST08, PG10]. Importantly, calcium signals capture not only the mean level of neural activity but also its variability [GK12]. We show that this dual encoding enables a novel solution to the E/I coordination problem. Specifically, the mean of the calcium signal regulates inhibitory plasticity, while the second moment of calcium fluctuations governs excitatory plasticity. Crucially, these processes act through distinct protein-synthesis pathways with limited resources, ensuring that excitatory and inhibitory synapses can co-adapt without mutual interference. This division of labor allows neurons to maintain a stable firing rate while preserving spiking variability—a hallmark of cortical activity in vivo across physiological conditions [SK93, MOS^+^16].

In the following sections, we develop this framework and analyze its consequences. Section 2.1 shows why classical homeostatic models with single-loop firing-rate feedback fail, leading to instability when both E and I synapses adapt concurrently. Section 2.2 introduces a multidimensional calcium-based model in which the calcium mean regulates inhibitory plasticity, while the second moment drives excitatory plasticity through distinct, resource-limited protein synthesis pathways. In Section 2.3, we demonstrate how such divergent encoding naturally arises from nonlinear dependencies of downstream signaling on the calcium trace. Section 2.3.1 establishes the link between calcium’s mean and firing rate, and between its variance and spiking irregularity. Section 2.3.2 derives the steady-state solutions of the closed-loop calcium–plasticity system, identifying conditions for stable convergence. Finally, Section 2.4 shows that this fixed point generates a balanced state, with synaptic weights scaling as 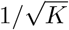 and maintaining realistic firing rates and irregular spiking. This ensures that E–I networks with diverse physiological parameters robustly settle into the balanced regime observed experimentally [MOS^+^16].

## 2 Results

### 2.1 The Failure of Single-Loop Feedback in E/I Co-Regulation

Classical firing-rate homeostasis models adjust synaptic weights via negative feedback on the output firing rate [DA05]. However, they fail to account for the simultaneous self-organization of both excitatory and inhibitory synapses.

To illustrate this limitation, consider a neuron receiving *K* excitatory and *K* inhibitory independent Poisson inputs at rates *λ*_*E*_ and *λ*_*I*_. Under the diffusion approximation [Kam07], the total input current can be modeled as

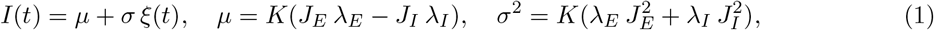

where *J*_*E*_ and *J*_*I*_ are the synaptic weights, and *ξ*(*t*) is standard Gaussian white noise with unit variance. The neuron’s steady-state firing rate is *r* = *f* (*µ, σ*), where *f* is its input–output transfer function.

A standard homeostatic rule for excitatory weights is

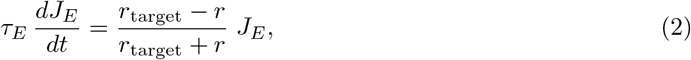

and analogously for inhibition,

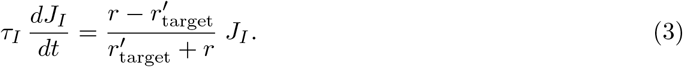

Each rule can independently stabilize its respective weight when the other is fixed. However, enforcing both simultaneously creates an overdetermined system. At equilibrium 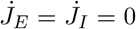, the fixed points imply

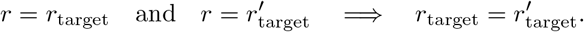

Under this condition, Eq. (1) constrains only the net input *J*_*E*_ *λ*_*E*_ − *J*_*I*_ *λ*_*I*_, allowing a continuum of weight ratios *J*_*E*_*/J*_*I*_. Any small perturbation or noise causes the weights to diverge rather than converge (see the Materials and Methods, section 4.1, for the stability analysis).

Maintaining 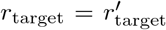 would require precise coordination between distinct molecular pathways, such as TNF*α*-mediated bidirectional scaling of excitatory and inhibitory synapses and BDNF-mediated upscaling of inhibition. However, these pathways operate independently and are activated under different activity regimes, making such coordination unlikely [SM06, PZS^+^10].

Previous models circumvent this failure by imposing ad hoc constraints, such as fixing one synapse type [AL93] or introducing external balancing mechanisms [LMA93]. Such assumptions undermine genuine self-organization and contradict experimental observations of concurrent, pathway-specific E/I plasticity.

This paradox highlights the need for a unified feedback mechanism that relies on a single post-synaptic activity signal while engaging independent control pathways for E and I synapses. In the following sections, we show that intracellular calcium dynamics fulfill these criteria by decomposing firing statistics into their mean rate and spiking variability components. Crucially, these calcium signals regulate excitatory and inhibitory plasticity through distinct, resource-limited protein-synthesis pathways, allowing both synapse types to adapt in parallel without interference.

### 2.2 Calcium-Dependent Mechanisms for Distinct E/I Co-Regulation

Recent work has shown that intracellular calcium dynamics serve as a shared sensor for activity-dependent synaptic regulation [PG10, IST08, GK12], while still permitting distinct downstream signaling pathways at E and I synapses that rely on separate, resource-limited protein-synthesis processes [Tur04].

We propose that **distinct calcium statistics—mean for inhibition and second moment for excitation—resolve the overdetermination problem by enabling two independent control pathways from a single activity signal**. Through their respective protein-synthesis constraints, this statistical decomposition enables the separate regulation of firing rate (via *J*_*I*_) and spiking irregularity (via *J*_*E*_), thereby circumventing the need for fine-tuned coordination between E and I pathways.

This framework provides a mechanistic explanation for experimental observations in which inhibitory synapses adapt inversely to the strength of excitation [Tur04], while overall spiking irregularity is preserved. To resemble this experimental observation, in our model we introduce two hypothetical calcium-dependent proteins, P_E_ and P_I_, that regulate E and I synapses through separate biochemical cascades. Both pathways follow Michaelis–Menten kinetics [JG11] and are driven by the somatic calcium concentration [Ca^2+^](*t*) [Ber98], but differ in their sensitivity to the temporal statistics of calcium fluctuations and in the limited pool of protein resources available for their synthesis.

#### 2.2.1 Inhibitory Synapse Regulation: Mean Calcium Dependence

We model inhibitory protein synthesis as a calcium-dependent enzymatic reaction. Let P_I_ be an intracellular protein that promotes the growth of inhibitory synapses. It is synthesized by an enzyme E_I_ that binds a calcium ion before catalyzing the production of P_I_, using a metabolic substrate X (e.g., amino acids or ATP). The relevant reactions are:

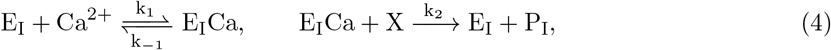

and P_I_ is degraded via:

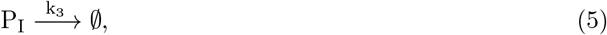

where degradation products are omitted for traceability as recycling is assumed to occur on faster timescales than P_I_ turnover. Furthermore, by assuming calcium and substrate X are abundant relative to E_I_, quasi-steady-state analysis yields:

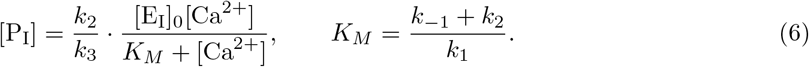

Under physiological conditions where [Ca^2+^] ≪ *K*_*M*_, this simplifies to:

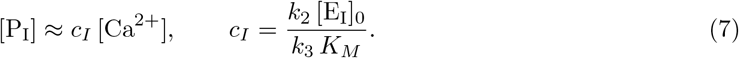

We model the change in inhibitory synaptic efficacy *J*_*I*_ as a slow adaptive process driven by deviations in [P_I_] from a homeostatic set point:

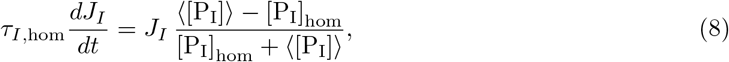

where ⟨·⟩ denotes temporal averaging over the protein turnover timescale *τ*_*I*,hom_. Here, [P_I_]_hom_ reflects the limited concentration of available protein to regulate inhibitory synapses.

Somatic calcium fluctuations occur on timescales ranging from tens to several hundred milliseconds, depending on cell type and buffering capacity [KS00]. In contrast, protein synthesis and degradation are much slower (*τ*_*I*,hom_ ∼ hours) [SSM^+^21, Tur08]. Thus, the averaged protein concentration integrates over many calcium transients:

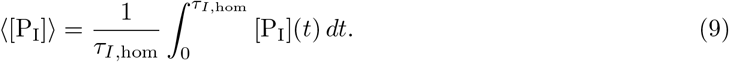

Substituting Eq. (7) and defining the homeostatic calcium set point as ⟨[Ca^2+^] _hom_ ⟩ ≡ [P_I_] _hom_ */c*_*I*_, corresponding to a limited regulatory portion in the neuron, thus, the plasticity rule becomes:

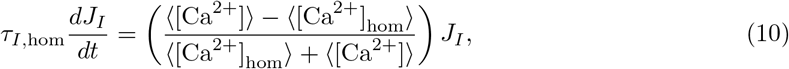

where ⟨[Ca^2+^]⟩ is the mean intracellular calcium concentration at steady state. This is consistent with classical negative feedback models, where firing rate *r* is approximately proportional to mean calcium level.

#### 2.2.2 Excitatory Synapse Regulation: Calcium Second Moment Dependence

Excitatory synaptic efficacy responds to activity fluctuations via **nonlinear calcium sensing**. We model the production of an excitatory-promoting protein P_E_ as driven by cooperative calcium binding to an enzyme E_E_ (e.g., following calmodulin-like kinetics [LYR12]), which uses a metabolic substrate Y:

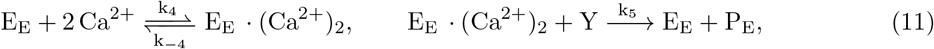

with degradation:

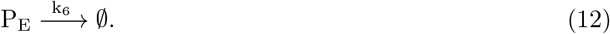

Assuming calcium and substrate Y are abundant relative to E_E_, and enforcing enzyme conservation [E_E_]_0_ = [E_E_] + [E_E_ · (Ca^2+^)_2_], we obtain the steady-state bound enzyme concentration:

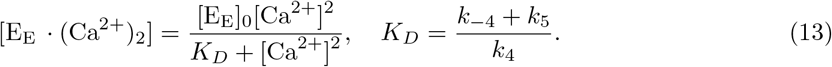

Balancing production and degradation yields:

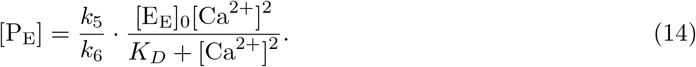

In the regime where [Ca^2+^]^2^ ≪ *K*_*D*_, this simplifies to:

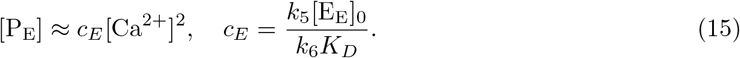

This quadratic dependence implies that P_E_ production is sensitive to calcium variability. Consider a calcium trace decomposed into a mean and zero-mean fluctuations:

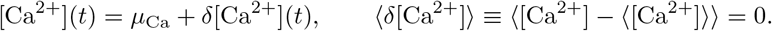

Then:

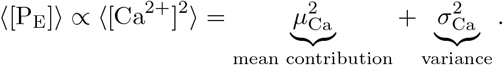

When inhibitory plasticity (Section 2.2.1) stabilizes *µ*_Ca_, the term ⟨[P_E_]⟩ becomes primarily sensitive to 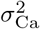, making it an effective *variance sensor*.

Accordingly, deviations from a homeostatic value of the second calcium moment set point regulate excitatory synaptic efficacy *J*_*E*_ as:

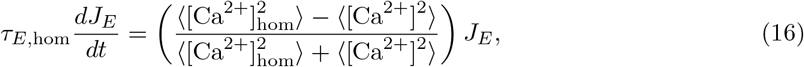

where 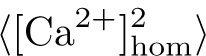 is the homeostatic target for P_E_ synthesis, and *τ*_*E*,hom_ denotes the protein turnover timescale (as in Eq. (10)), where is usually *τ*_*E*,hom_ *> τ*_*I*,hom_ [Fro15, Tur17]. This cooperative calcium binding mechanism (*n* = 2) is consistent with calmodulin/CaMKII signaling pathways [LYR12]. In the next section, we show that the regulation of calcium second moment enables the stabilization of spiking irregularity.

### 2.3 Calcium Homeostasis Framework for General Spiking Neurons

Intracellular calcium serves as a key biochemical integrator of spike train statistics, mediating activity-dependent synaptic plasticity. To understand how calcium dynamics can implement closed-loop homeostasis of excitatory and inhibitory inputs, we establish a general theoretical framework that links spike timing statistics to intracellular calcium concentration.

This framework applies to any spiking neuron model with stationary statistics. It closes the loop initiated in Sections 2.2.1 and 2.2.2, where we derived synaptic homeostatic rules based on calcium moments. By expressing calcium statistics in terms of the spike train, we obtain a self-consistent description of E/I co-regulation.

#### 2.3.1 Calcium Dynamics Driven by Spiking Statistics

Consider a neuron emitting spikes at times *t*_*n*_, modeled as a stationary renewal process with mean rate *r* = ⟨*S*(*t*)⟩, where *S*(*t*) Σ = *δ*(*t* − *t*_*n*_). The Inter-Spike Interval (ISI) distribution is given by *p*(Δ*t*). Intracellular calcium concentration evolves via hybrid dynamics: between spikes,

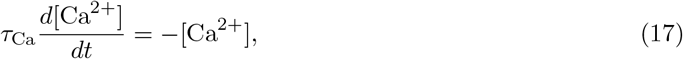

and at spike times *t*_*n*_, it jumps according to:

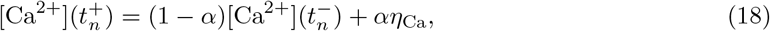

where *τ*_Ca_ is the calcium decay constant, *η*_Ca_ is the unitary calcium influx per spike, and *α* ∈ (0, 1] models fast buffering [MZS04, SJ05].

Using the Laplace transform of the ISI distribution,

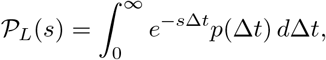

we derive closed-form expressions for the calcium moments (see Materials and Methods Section 4.2 for derivations):

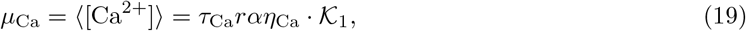

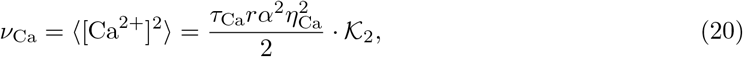

where the dimensionless factors 𝒦_1_ and 𝒦_2_ are defined as:

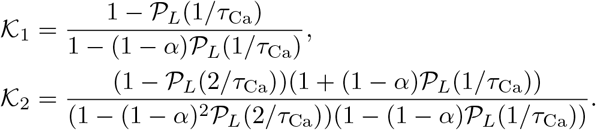

In Materials and Methods (Theorem 2), we proved that, for any ISI family ordered in convex order and under a mild balanced-buffering condition (satisfied for physiologically small *α*), the two dimensionless kernels

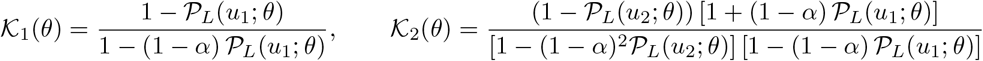

satisfy

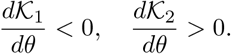

Thus, increases in ISI variability (higher convex-order ISIs) strictly *decrease* 𝒦_1_ and *increase* 𝒦_2_. We refer to this bidirectional sensitivity as **divergent encoding** of somatic calcium signals.

As a concrete example, let us consider the Gamma distribution for the ISI,

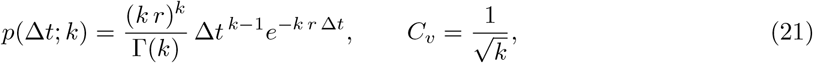

so that larger *C*_*v*_ corresponds to smaller *k*. Its Laplace transform at *s* = *u*_*i*_ is

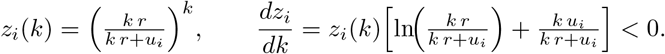

Since 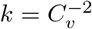 implies

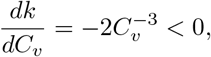

the sign of 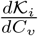 is opposite to that of 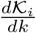. Then one computes

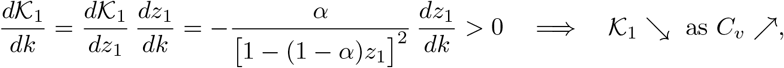

and, under the same buffering condition on *α*,

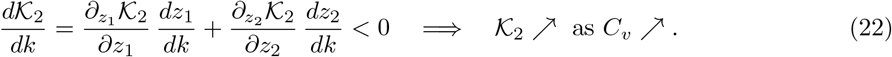

Thus for physiologically relevant buffering (*α* small), Gamma-distributed ISIs of equal mean but increasing *C*_*v*_ produce exactly opposite trends in 𝒦_1_ and 𝒦_2_, and hence in the calcium mean *µ*_Ca_ and second moment *ν*_Ca_. This derivation shows that the Gamma distributed ISI follows the **divergent encoding** of somatic calcium signals, as proven in Theorem 2 (Materials and Methods, Section 4.3).

To illustrate the divergent-encoding result of Theorem 2 (and the derivation above), we plot how calcium statistics vary with spike-train irregularity using Gamma-renewal ISIs at a fixed firing rate (Eq. (21)). Figure 1A shows that the mean calcium concentration *µ*_Ca_ decreases slightly as the coefficient of variation *C*_*v*_ increases, whereas Figure 1B shows that the second moment *ν*_Ca_ rises sharply. This divergent dependence confirms that somatic calcium can simultaneously encode firing rate and irregularity via distinct statistical moments, yielding separable signals for modulating the strength of inhibitory and excitatory synapses.

**Figure 1:**
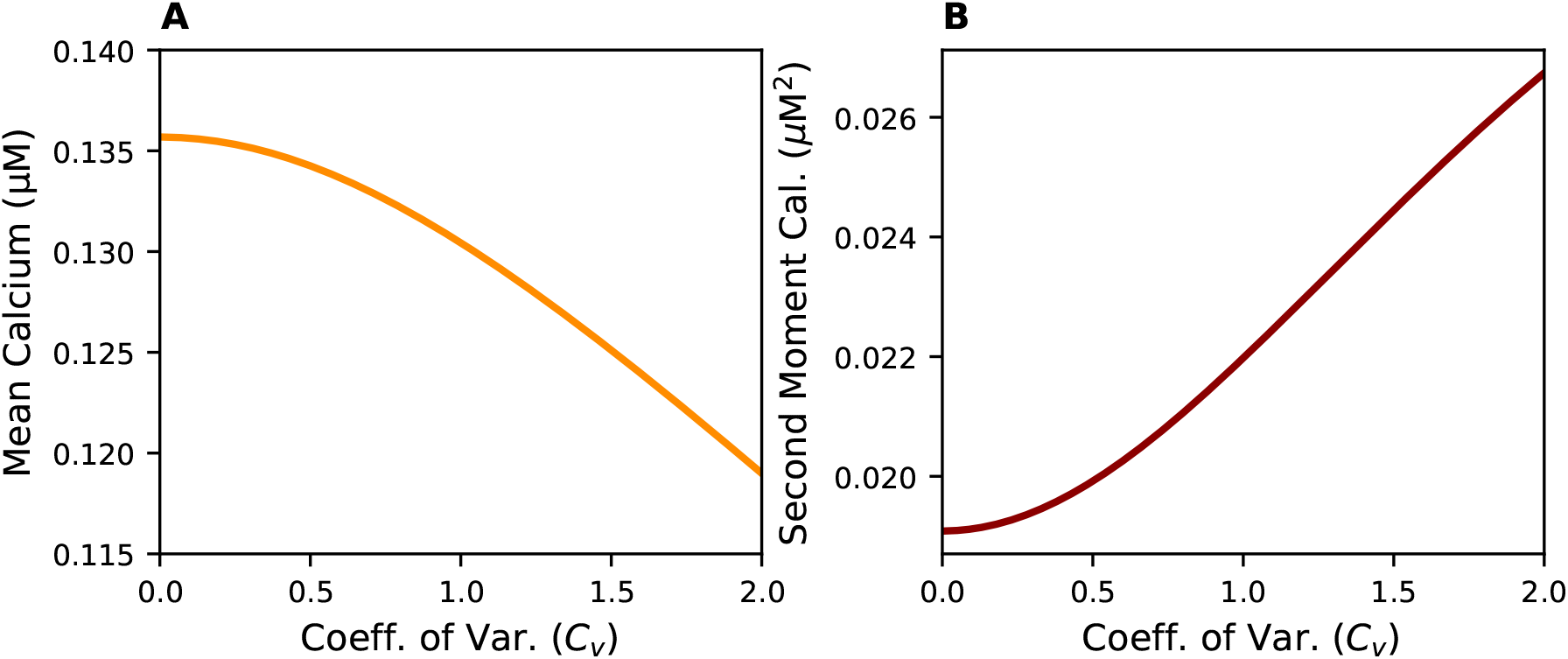
Calcium moments diverge with increasing spiking irregularity. (A) Mean calcium concentration *µ*_Ca_ as a function of the coefficient of variation *C*_*v*_ for a Gamma renewal process at fixed firing rate *r* = 5 Hz. (B) Corresponding second moment *ν*_Ca_. Simulations confirm theoretical predictions from Eqs. 19–20. Parameters: *α* = 0.1, *η*_Ca_ = 1 *µ*M, *τ*_Ca_ = 300 ms. While the mean decreases modestly with *C*_*v*_, the second moment increases strongly, supporting the divergent encoding of rate and variability by the mean and second moment of calcium fluctuation.

#### 2.3.2 Closed-Loop Homeostatic Dynamics

The closed-loop system emerges by coupling calcium-based synaptic plasticity rules to spike-driven input statistics. Specifically, for the input structure as in Eq. (1), the mean and variance of synaptic input are determined by presynaptic firing rates and synaptic weights, which closes the feedback loop via calcium dynamics. Input statistics, together with neuronal spiking dynamics, determine ISI density, which specifies the calcium moments (Eqs. 19 and 20). Substituting the calcium moments into the plasticity rules (Eqs. 10 and 16) yields a self-contained dynamical system for synaptic weights:

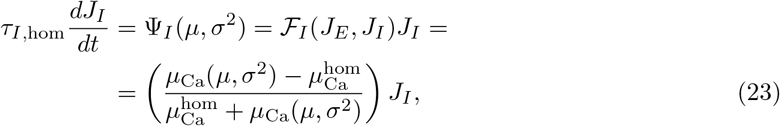

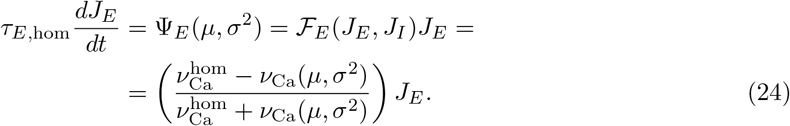

Here, the calcium moments *µ*_Ca_ and *ν*_Ca_ depend implicitly on the synaptic weights via the input statistics defined in Eq. (1). This creates a feedback cascade in which synaptic weights modulate neural input, shaping spiking activity and intracellular calcium, which ultimately feed back to drive synaptic regulation.

The system reaches equilibrium when the calcium moments match their homeostatic targets:

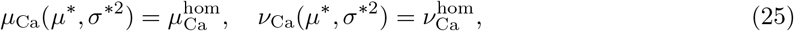

corresponding to steady-state input statistics

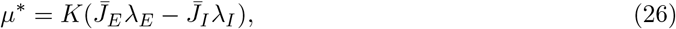

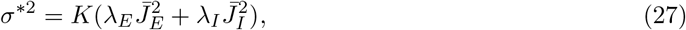

where 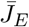 and 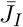 are the equilibrium synaptic weights. Because *µ*_Ca_ and *ν*_Ca_ encode complementary aspects of spiking due to Theorem 2, these equilibrium conditions are jointly solvable across a wide range of ISI distributions with finite mean and variance observed in spiking neurons [Tuc05]. The physiological ISI density generally results from neuronal spiking, where:

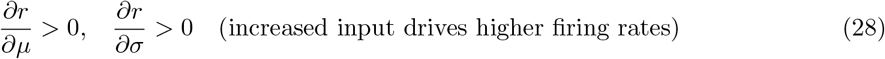

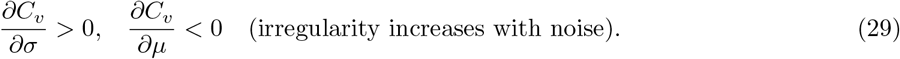

Standard spiking models meet these conditions in in vivo-like fluctuation-driven regimes, including leaky and exponential integrate-and-fire neurons, as well as noisy threshold models.

To assess the dynamic stability of the solution, we linearize the system around the fixed point. The Jacobian, shaped by the divergent calcium sensitivities as in Theorem 2, satisfies tr(*J*) *<* 0 and det(*J*) *>* 0 under conditions in Eq. (29), implying asymptotic convergence to a stable node under broad physiological conditions (for all details of stability analysis see Theorem 3 as it is discussed in Materials and Methods in Section 4.4).

To concretely illustrate the fixed-point structure and verify the stability predicted by Theorem 3 in Materials and Methods (Section 4.4), we analyzed the closed-loop dynamics in a perfect integrate-and-fire (PIF) neuron model, chosen for its analytical tractability. The PIF model yields closed-form expressions for the Laplace transform of the ISI distribution, the mean firing rate, and the coefficient of variation (*C*_*v*_) as functions of the input statistics (*µ, σ*), enabling a complete analytical characterization of the closed-loop dynamics (see Section 4.5.1).

Figure 2 illustrates the nullclines of the homeostatic dynamics derived from the calcium-based feedback model. Fig. 2A plots the nullclines 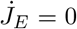 and 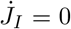 in the (*µ, σ*) space of synaptic input statistics. The intersection of these curves marks a unique fixed point of the dynamics where both the calcium mean and variance match their respective homeostatic setpoints, confirming that the system admits a consistent and jointly solvable equilibrium. In Fig. 2B, we project these nullclines into the space of observable firing statistics (*r, C*_*v*_) using analytical expressions specific to the PIF model as it is given in Section 4.5.1. The mapped fixed point aligns precisely with the target values *r*^*∗*^ and 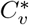 implied by the homeostatic constraints, indicating that the calcium-based plasticity mechanism successfully drives the neuron to a homeostatic target statistics.

**Figure 2:**
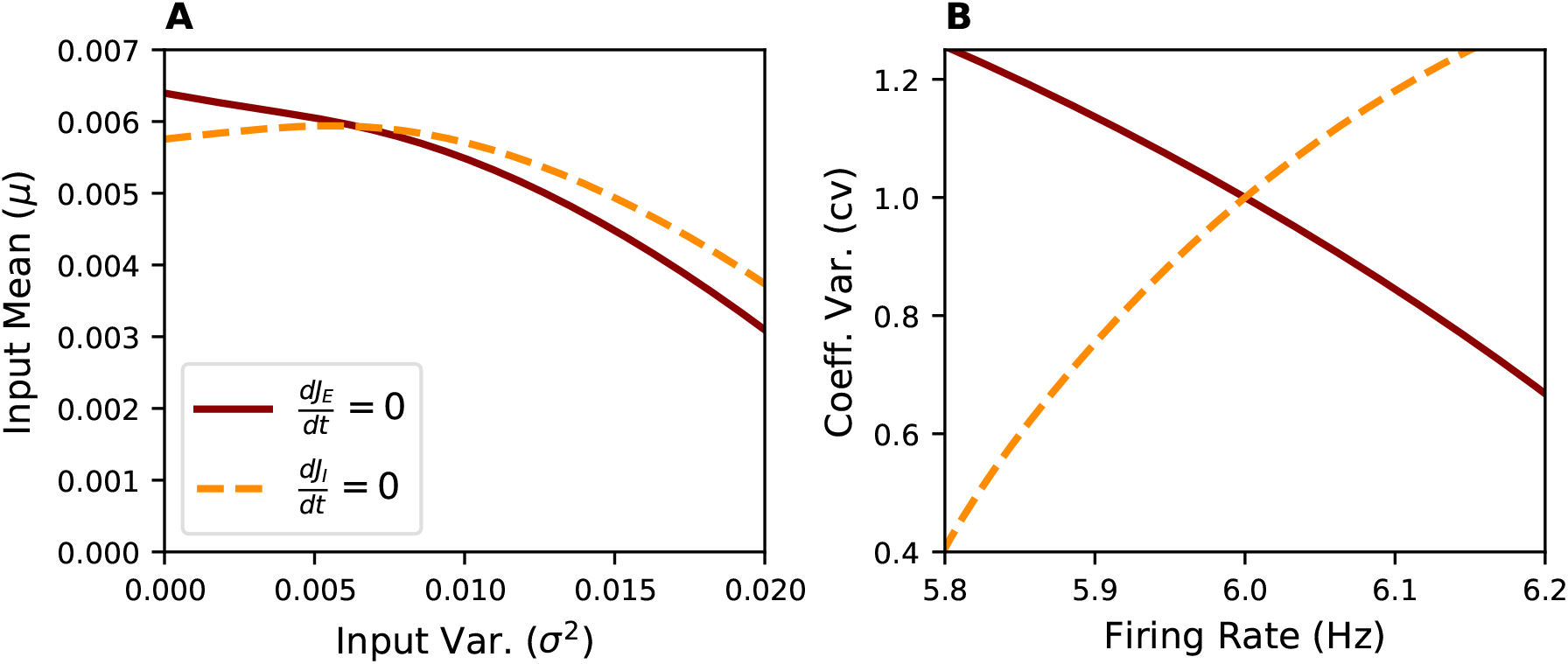
Nullclines of the closed-loop dynamics in the PIF model. **(A)** Nullclines 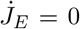 and 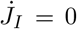 in the space of input statistics (*µ, σ*). Their intersection marks the unique fixed point where both calcium homeostasis constraints are satisfied. **(B)** Same nullclines as in (A) mapped into activity space (*r, C*_*v*_) using analytical PIF expressions in Materials and Methods (Neuronal parameters: *v*_*th*_ = 10, *v*_*r*_ = 0 and *v*_*B*_ = −1). The fixed point aligns with the target activity levels *r*^*∗*^ and 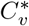.

Together, these results establish that the closed-loop system, driven by calcium’s dual encoding of mean and second moment, converges robustly to a stable fixed point. This mechanism enables autonomous regulation of both postsynaptic firing rate and spiking irregularity.

### 2.4 Emergence of the Balanced State

We now analyze how the calcium-based E-I feedback loop shapes synaptic efficacies at the stable fixed point of Eqs. (23)–(24). Specifically, we examine how the homeostatic constraints on calcium moments translate into a scaling relationship between synaptic weights and the number of presynaptic inputs *K*. We demonstrate that the intracellular calcium dynamics impose a principled scaling law on synaptic strengths as the number of synapses, *K*, increases. Importantly, this scaling ensures that single-neuron input fluctuations remain of order one, supporting irregular spiking. When generalized to a recurrent network, the regulation of calcium multi-dimensional signal gives rise to the classical balanced state [vVS98], characterized by dynamically balanced excitation and inhibition, strong temporal fluctuations at the population level, and broad heterogeneity in individual firing rates [RBH^+^11, vVS98].

#### 2.4.1 Single-Neuron Synaptic Scaling under Calcium Homeostasis

As it is introduced in Eq. (1), we consider a single neuron receiving *K* excitatory and *K* inhibitory Poisson inputs at fixed rates *λ*_*E*_ and *λ*_*I*_, respectively. Under the diffusion approximation, the stability and existence of the fixed point dynamics for our homeostatic model (Section 2.3.2), the mean and variance of the total synaptic input at equilibrium are given by Eqs. (26) and (27). Solving Eqs. (26)–(27) for equilibrium weights yields

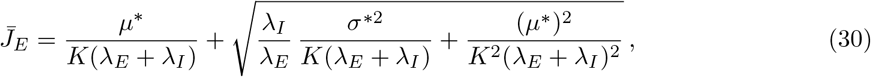

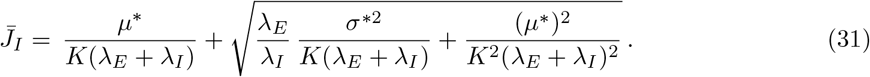

In the large-*K* limit, the square-root terms dominate the *O*(1*/K*) offsets, yielding the asymptotic behavior

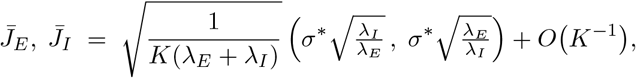

so that 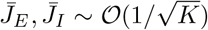. This leads to the emergence of van Vreeswijk [vVS98] scaling solution to achieve dynamic balance input,

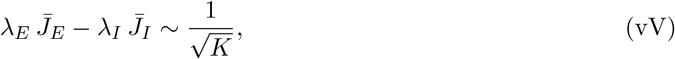

where the net input mean and variance remain finite as *K* → ∞, i.e., *µ*^*∗*^ ∼ 𝒪(1), ∼ *σ*^*∗*2^ 𝒪 (1) [vVS96, vVS98, vVS05].

To verify the equilibrium solutions scaling for weights given in Eqs. (30) and (31), we integrate weight dynamics in Eqs. (23) and (24) for a PIF neuron to achieve desired homostatic *r*^*∗*^ and 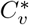 for the fixed-point as in Figure 2B while varing number of input *K*. Fig 3A illustrates the equilibrium synaptic weights 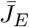 (red solid line) and 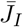 (orange dashed line) as a function of increasing input number *K* under calcium homeostasis. Both weights decrease with *K*, reflecting the predicted weakening of individual synapses as the number of inputs grows. Fig 3B further plots the empirically estimated scaling exponent *γ*_0_ (defined by 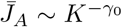 for *A* ∈ {*E, I*}) for each weight, derived from the results in Panel A. As *K* increases, both exponents converge to the theoretical prediction *γ*_0_ = 0.5, confirming the asymptotic 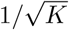 scaling due to homeostasis rules in Eq. (26)–(27) at equilibrium.

**Figure 3:**
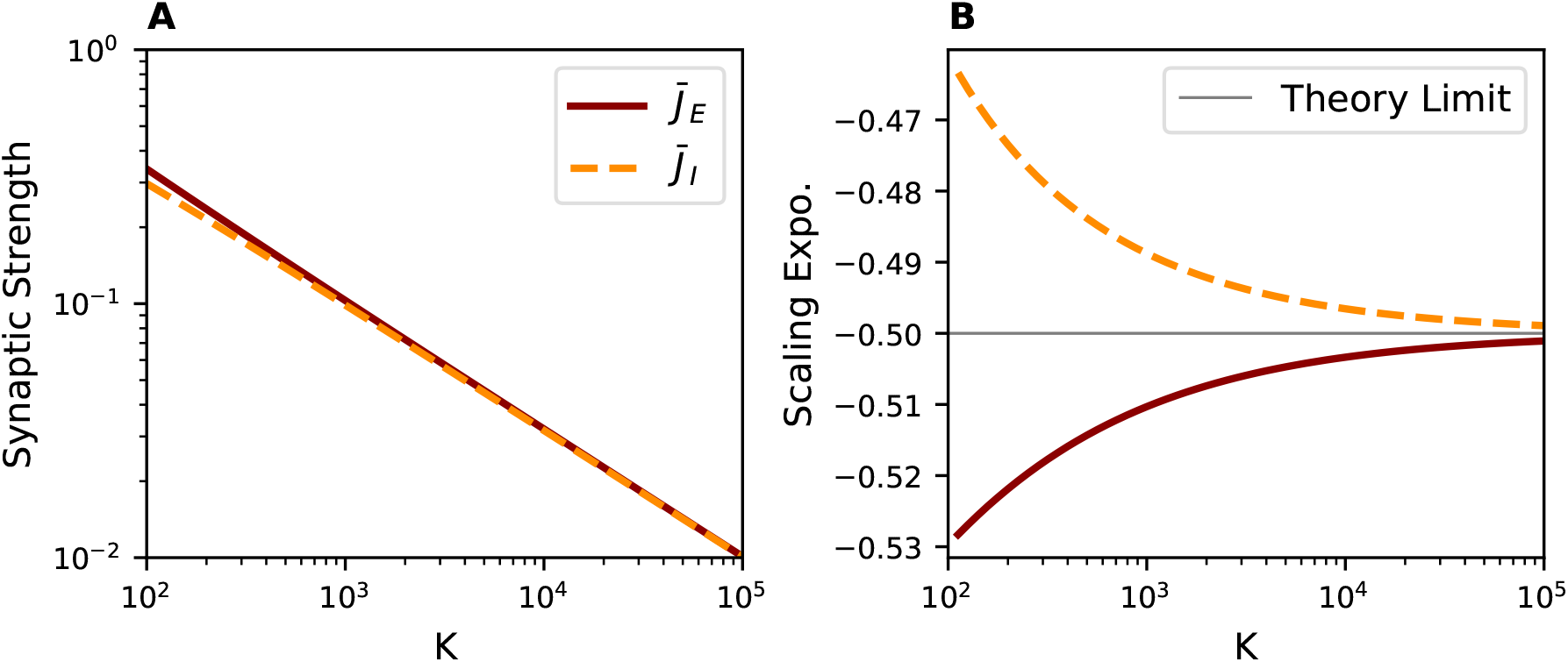
Emergence of the balanced state under E-I homeostasis. **(A)** Equilibrium excitatory (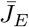, red solid line) and inhibitory (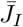, orange dashed line) synaptic weights as a function of the number of presynaptic inputs *K* in the perfect integrate-and-fire model. Both weights decrease with *K*, following a 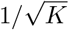 scaling predicted by calcium-based homeostasis. **(B)** Empirically estimated scaling exponent *γ*_*0*_ (defined by 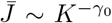) for each weight, showing convergence to the theoretical prediction *γ*_*0*_ = 0.5 as *K* increases. Analytical predictions are based on closed-form expressions for calcium moments for the perfect integrate-and-fire neuron model, as derived in the *Materials and Methods* Section 4.5.1, with parameters *v*_th_ = 10, *v*_*r*_ = 0, and *v*_*B*_ = −1.

#### 2.4.2 Excitatory–Inhibitory Balance on the Network Level

In large recurrent networks, heterogeneity in connectivity generates a broad distribution of input statistics and firing rates. To investigate how calcium homeostasis enforces balance in such a network, we embed our calcium-dependent plasticity rules into an E–I circuit with sparse Erdős–Rényi connectivity (1 ≪ *K* ≪ *N*). In the large-*K* limit, each neuron *i* in population *A* ∈ {*E, I*} receives *k*_*iAB*_ inputs from population *B* ∈ {*E, I, X*}, where *k*_*iAB*_ can be approximated by a Gaussian distribution with mean *K* and variance *K*: *k*_*iAB*_ ∼ 𝒩 (*K, K*). Here, *X* denotes an external excitatory Poisson drive at a fixed rate *r*_*X*_. Under the diffusion approximation (see Materials and Methods, Sec. 4.5.3), similar to Eq. 1, the synaptic input to neuron *i* in *A* is

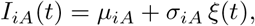

with

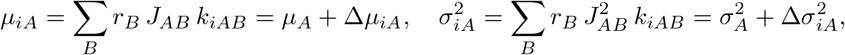

and population-mean statistics

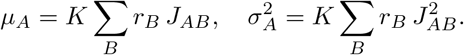

Here, Δ*µ*_*iA*_ and 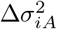 are zero-mean Gaussian random variables with variances

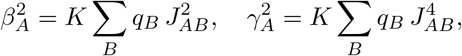

where

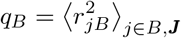

is the second moment of the firing-rate distribution in population *B*, and ⟨·⟩_***J***_ denotes averaging over connectivity realizations. Hence, 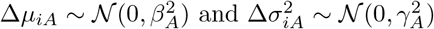.

Each neuron’s firing rate is given by 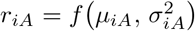. Thus, the population-averaged firing rate *r*_*A*_ and its second moment *q*_*A*_ satisfy

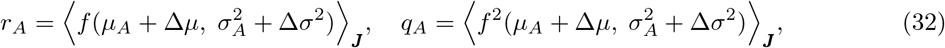

where the averages denote expectation over (Δ*µ*, Δ*σ*^2^) drawn from the bivariate Gaussian (Methods, Sec. 4.5.3).

Homeostatic plasticity updates the mean synaptic strengths according to

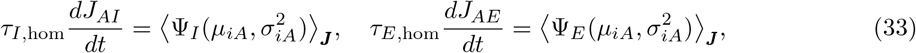

where Ψ_*I*_ and Ψ_*E*_ implement the calcium-based inhibitory and excitatory update rules using the synaptic input statistics of the postsynaptic population *A* (following Eqs. (10) and (16)). Note that the synaptic update rule from the external population *X* follows the excitatory rules. Both depend on the homeostatic target point set for each population *A*, defined using the asymptotic balanced-network statistics:

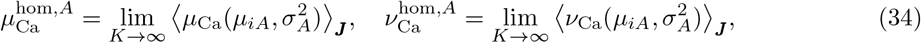

where averaging follows the statistics of the balanced state in the thermodynamic limit (*K* → ∞), using the self-consistent solution of the balanced van Vreewsijk equation for rate 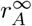, and quenched disorder 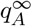 (see Materials and methods in section 4.5.4).

To allow the finite-*K* network to converge to a self-consistent equilibrium, the homeostatic targets continuously adapt toward the population average instantaneous calcium statistics:

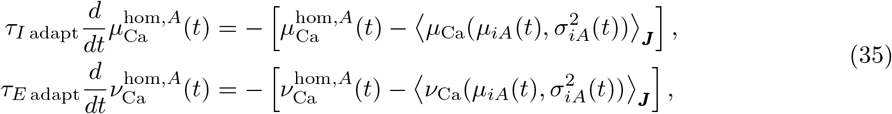

where *τ*_*A* adapt_ sets the adaptation timescale, and it is assumed to be 10 times bigger than *τ*_*A* hom_. This continuous alignment ensures that the network maintains a functional equilibrium despite the *finite* resources available in the system.

To investigate the emergence of balanced network dynamics under calcium-based homeostasis, we numerically solve the self-consistent mean-field equations including quenched correlation structure (see Materials and Methods Sec. 4.5.3) by integrating the synaptic weights according to Eq. (33), with dynamic homeostatic targets governed by Eq (35). We assume a timescale separation in which neuronal firing rates and input fluctuations equilibrate rapidly relative to synaptic changes in Eq. (33). Neuronal spiking rates and ISI density are given using the Leaky Integrate-and-Fire (LIF) model (see Materials and Methods, Section 4.5.2).

Figure 4 illustrates that calcium-dependent E-I plasticity robustly drives the network toward a fixed point exhibiting the hallmark features of the E-I balanced state. Panel A shows that as the in-degree *K* increases, both excitatory and inhibitory synaptic weights follow the characteristic 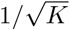 scaling law (Fig. 4A). At the same time, the population-averaged firing rates *r*_*E*_ and *r*_*I*_ converge to their theoretical large-*K* predictions, i.e. 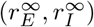 (Fig. 4B). Moreover, calcium homeostasis stabilizes both the temporal variance of synaptic input 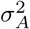 (Fig. 4C) and the quenched disorder variance 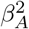 (Fig. 4D), in agreement with the analytically predicted statistics of a balanced state (see Materials and Methods section 4.5.3). Together, these results demonstrate that the proposed homeostatic rules generalize the mechanism of calcium-mediated resource allocation for synaptic regulation of recurrent circuits, enabling the emergence of network-wide balance that preserves temporal fluctuation and heterogeneity due to quenched disorder.

**Figure 4:**
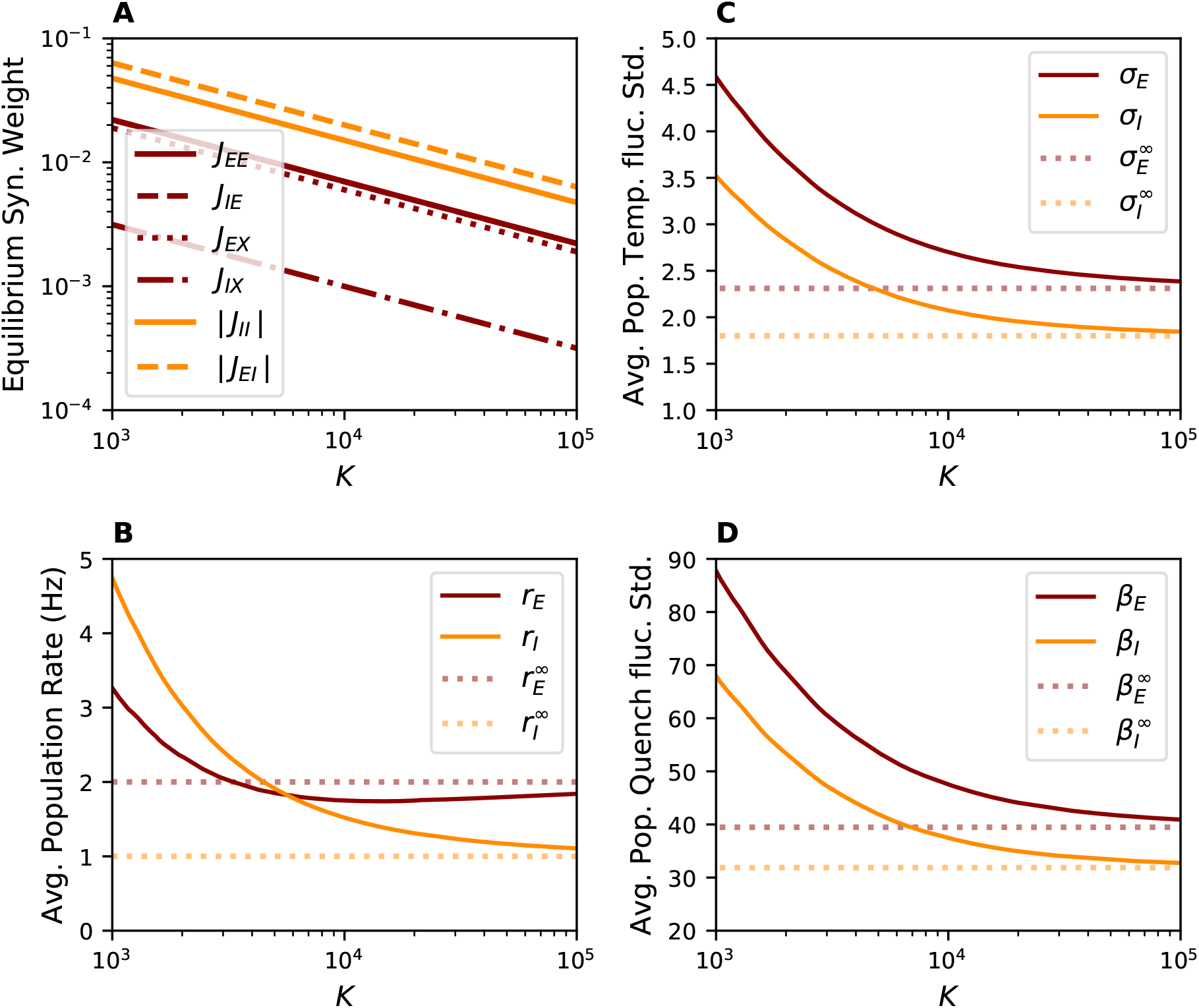
Emergence of balanced state in E-I network dynamics via calcium homeostasis. Self-consistent mean-field solutions for a recurrent *E*–*I* network under calcium-based synaptic plasticity. (**A**) The excitatory (*J*_*AE*_ and *J*_*AX*_, red) and inhibitory (*J*_*AI*_, orange) synaptic weights scale as 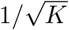 with increasing average in-degree *K* in Erdős–Rényi networks, consistent with balanced-state theory, where *A* ∈ {*E, I*} is the postsynaptic population. (**B**) Population-averaged firing rates *r*_*E*_ and *r*_*I*_ converge to their predicted values in the large-*K* limit. (**C**) Temporal variance of synaptic input fluctuations 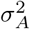, and **D**) The variance due to quenched disorder, 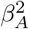. They match the statistics predicted for a balanced state mean-field theory. Together, these results confirm that calcium-dependent homeostasis stabilizes both the population rate and temporal fluctuation variance while maintaining heterogeneous firing across the network. The underlying neuron model used to solve the mean-field equations is a leaky integrate-and-fire (LIF) neuron with parameters *τ*_*m*_ = 10 ms, *v*_th_ = 1.0, *v*_*r*_ = 0.0, and *τ*_ref_ = 2 ms.

## 3 Discussion

Neurons in the cortex must continuously adapt to changing inputs while retaining stable function. Our work demonstrates that intracellular calcium dynamics—long recognized as a key integrator of synaptic activity—can provide two independent feedback channels that support the emergence of balanced cortical networks. By mapping the mean calcium signal to inhibitory synaptic homeostatic mechanisms and the second moment of calcium fluctuations to excitatory ones, we show how distinct calcium-binding kinetics in molecular cascades [Ber98], together with resource-limited protein-synthesis pathways, can encode orthogonal feedback signals. This division of labor enables robust co-regulation of E and I synapses without the need for fine-tuned parameters or external control mechanisms.

We analytically characterized the closed-loop calcium dynamics driven by spike trains with diverse interspike interval statistics (Section 2.3). Our analysis revealed that the calcium mean increases monotonically with firing rate, while the calcium second moment is more sensitive to spike-train irregularity and relatively insensitive to rate (Section 2.3.1). These complementary sensitivities allow inhibitory homeostasis to stabilize mean firing rates, while excitatory synaptic regulation preserves temporal variability essential for neural coding [SK93, MOS^+^16]. We further prove that our dual calcium-dependent E-I homeostatic rules form a stable fixed point under general neuronal input–output properties (Section 2.3.2). We demonstrate this fixed-point stability in the PIF neuron model (Figure 2). Thus, calcium-based feedback provides a biologically plausible mechanism for the co-regulation of rate and spiking irregularity without disrupting ongoing dynamics.

Experimental evidence supports the neuronal plausibility of our framework in which distinct statistics of the calcium signal drive excitatory and inhibitory plasticity through separate molecular pathways. Sustained elevations in intracellular calcium preferentially activate calcineurin–NFAT and TNF*α*-mediated cascades that promote inhibitory scaling [SM06, PZS^+^10]. By contrast, transient and high-variance calcium signals engage CaMKII and CaMKIV pathways that selectively modulate excitatory plasticity [IST08]. These pathways rely on resource-limited protein synthesis, yet their distinct sensitivities to the temporal statistics of calcium allow inhibition to be tuned by mean activity levels and excitation by fluctuations, thereby supporting stable co-adaptation without mutual interference.

Irregular spiking and fluctuation-driven regimes are widespread across species and are essential for cortical function [MOS^+^16]. The balanced state theory has been used to explain the irregular spiking of cortical circuits, without the fine-tuning of parameters [vVS98, vVS96]. The hallmark of the theory is based on the hypothesis that synaptic weights scale as 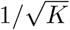, which leads to order one temporal fluctuations in synaptic input while the leading-order mean input dynamically cancels. A recent experimental observation in cortical cultures supports the notion that synaptic scaling follows 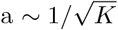 dependence on in-degree [BDR16], which matches the theoretical predictions.

The biological basis for the presence of balanced state spiking has been unknown. The calcium-dependent E-I homeostatic dual signaling remarkably allows for the emergence of a balanced state in various physiological conditions, as it naturally maintains 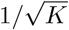 synaptic scaling law. We show that the stable fixed-point of the homeostatic dynamics corresponds to a balanced solution on the signal neuron level (Figure 3). We further show that embedded in sparsely connected recurrent networks, our calcium-dependent E-I feedback rules self-organize synaptic strengths to match the classical 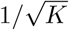 scaling of balanced networks [vVS98, vVS96], while preserving firing-rate heterogeneity and high variability across neurons [RBH^+^11]. This result underscores how local, cell-autonomous calcium feedback circuits can give rise to emergent, biologically plausible network-level balance.

Our framework also motivates direct experimental tests. Since our theory predicts that the calcium mean drives inhibitory scaling, in contrast, excitatory scaling depends on calcium variance. Therefore, optogenetic or pharmacological manipulation of the temporal statistics of synaptic input without altering the mean rate should differentially affect E/I balance. Specifically, increasing input irregularity should preferentially trigger excitatory downscaling, while smooth tonic input should activate inhibitory homeostasis. These predictions can be tested using calcium imaging combined with dynamic clamp or patterned optogenetics. In particular, our theory makes quantitative predictions for the effect of continuous optical stimulation in cultured networks, similar to the setting in [BDR16]. In a condition that a population is chronically driven by a weak optical current *I*_opt_, the recurrent weights self-organize such that both the mean and variance of calcium match their homeostatic targets, thus input statistics are given by

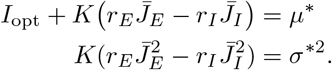

In this regime, the average firing rate and spike-train irregularity (*C*_*v*_) remain unchanged despite ongoing stimulation. For stronger stimulation, 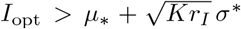, however the network cannot maintain balanced excitation. Therefore, the homeostatic rules compensate by driving excitatory weights to zero (*J*_*E*_ → 0), and only the calcium mean is homeostatically regulated. In this high-input regime, both firing rate and *C*_*v*_ increase with *I*_opt_. This prediction could be tested using chronic optical stimulation and electrophysiological recordings in vitro.

While our theoretical analysis employs a diffusion approximation to characterize calcium dynamics, this assumption may be challenged in small networks where finite-size effects introduce weak temporal correlations [Far25]. Such deviations can alter the precise scaling of synaptic weights, which is only critical for defining the thermodynamic limit of the system. Notably, the core mechanism of divergent calcium encoding remains intact: the first and second moments of calcium fluctuations reliably capture firing rate and spiking irregularity, even under non-idealized conditions. Consequently, homeostatic dynamics stabilize activity across a wide range of network sizes, consistent with in vivo cortical recordings. Instability could, in principle, arise if a finite network were to push toward the theoretical scaling limit. The adaptive rule in Eq. (35) acts as a form of meta-learning that flexibly allocates limited protein-synthesis resources. This ensures that the system settles into a stable equilibrium appropriate for its size, rather than encountering numerical instabilities toward an unphysiological theoretical limit.

Several extensions of our work warrant further exploration. While our model assumes current-based synaptic inputs, biological neurons operate under conductance-based synaptic drive, where input currents depend nonlinearly on membrane potential [DRP03]. This voltage dependence alters the relationship between synaptic input and calcium dynamics, potentially modifying the mapping between input statistics and homeostatic signals. It remains an open question whether dendritic nonlinearities in vivo-like conditions, as suggested in [DRP03, vVF21], could preserve the calcium moments that underpin our homeostatic rules. Exploring how calcium-dependent E-I homeostasis functions in conductance-driven regimes may reveal essential constraints on the tuning of plasticity mechanisms to the biophysical properties of synaptic integration.

Additionally, our model treats neurons as homogeneous and untuned, governed by the mean-field assumptions, which raises the question of how functional selectivity—such as orientation tuning in the sensory cortex—is preserved under global homeostatic rules. One possibility is that dendritic compartmentalization of calcium signals enables branch-specific homeostasis, allowing synapse-specific regulation without disrupting circuit-level selectivity [YMH00]. Interneuron diversity (e.g., PV^+^, SST^+^, VIP^+^) may also introduce multiple inhibitory control pathways that selectively regulate subnetworks [PXH^+^13]. Future work should investigate how such structural and molecular specializations allow calcium-based specific E-I feedback to coexist with stable functional tuning.

Finally, although our framework focuses on non-associative synaptic scaling, it may naturally complement Hebbian plasticity mechanisms that drive fine-scale connectivity resulting from ongoing learning within the system. By stabilizing global excitability and suppressing runaway dynamics, calcium-based homeostasis could create a permissive background in which associative plasticity sculpts stimulus-specific responses [LGR19]. In this scenario, global calcium signals maintain network stability, while local, activity-dependent changes—e.g., through NMDA receptor activation or dendritic coincidence detection—enable selective learning. Such a dual mechanism may explain how networks sustain sparse, robust representations despite ongoing synaptic turnover and plasticity [KHB17, CAZ^+^25].

In conclusion, by harnessing the multidimensional nature of intracellular calcium dynamics, we provide a unified and biologically grounded framework for excitatory–inhibitory homeostasis that links molecular signaling to emergent network structure. Our theory shows how stability and irregularity—two hallmarks of cortical activity—can arise from self-organizing plasticity without external tuning. This calcium-based E-I mechanism offers new insight into how neural circuits remain robust to perturbations while retaining the flexibility to adapt, learn, and compute in ever-changing environments.

## 4 Materials and Methods

### 4.1 Instability of Single-Loop E/I Feedback

We provide a detailed analysis of the non-existence of a joint fixed point when 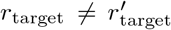 in models that only use the firing rate feedback to co-regulate E and I synaptic input. The homeostatic rules

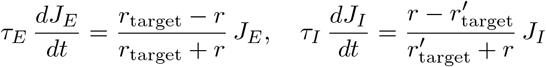

require at equilibrium 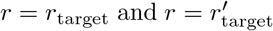 simultaneously. If 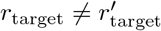, there is no solution 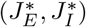 for which both derivatives vanish. Thus, no static operating point exists unless the two targets are equal, implying an unrealistically precise matching of E and I set-points.

When 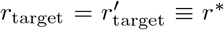, the system admits a one-dimensional continuum of equilibria. These satisfy

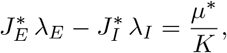

where *µ*^*∗*^ is fixed by *f* (*µ*^*∗*^, *σ*^*∗*^) = *r*^*∗*^, and *σ*^*∗*^ depends also on 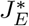 and 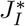 (through 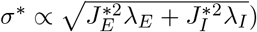. To assess stability, we linearize Eqs. (2)–(3) around an arbitrary point on this manifold. Denote small perturbations 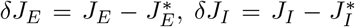, and write *δr* = *β* [*λ*_*E*_ *δJ*_*E*_ − *λ*_*I*_ *δJ*_*I*_] *K*, where 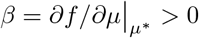 under physiological conditions. The linearized dynamics are

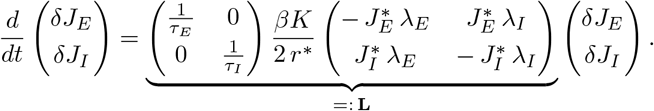

One computes det **L** = 0 and tr **L** *<* 0. Hence, the Jacobian has eigenvalues {0, tr **L**}, with one neutral direction (along the equilibrium manifold) and one stable direction (transverse to the manifold). In practice, any fluctuation along the neutral manifold (e.g., due to noise or parameter mismatch) is unopposed and leads to unbounded drift of *J*_*E*_ and *J*_*I*_.

When 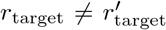, the absence of a fixed point implies the system cannot reach equilibrium. Near the manifold for equal targets, the dynamics develop a constant drift velocity proportional to 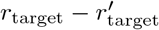, causing sustained changes in synaptic strengths. This drift can be approximated as

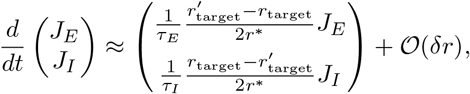

leading to unbounded evolution of synaptic weights.

### 4.2 Calcium Statistics for Renewal Processes

#### Theorem 1

(Calcium Statistics for Renewal Processes with Buffering). *For a stationary renewal spike train with ISI density p*(Δ*t*), *firing rate r* = ⟨*S*(*t*)⟩, *and Laplace transform* 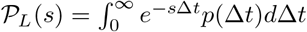, *consider the somatic calcium concentration C*(*t*) *has the following dynamics*

- *Between spikes:* 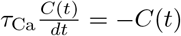*(exponential decay)*
- *At each spike time t*_*n*_: 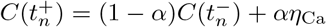 *(instantaneous buffering)*

*The steady-state calcium mean and second moment are*

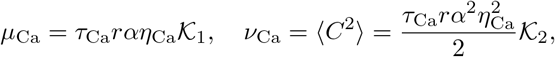

*where*

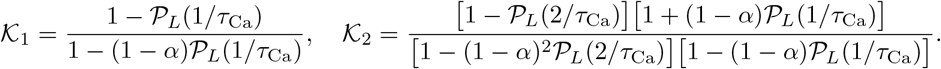

*Proof*. We derive the mean and second moment for this calcium dynamic.

#### Mean Calcium Concentration

Let 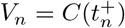 be the calcium concentration just after the *n*-th spike. The recurrence relation is:

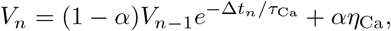

where Δ*t*_*n*_ = *t*_*n*_ − *t*_*n−*1_ is the *n*-th ISI. By stationarity and renewal property, ⟨*V*_*n*_⟩ = ⟨*V*_*n−*1_⟩ = ⟨*V* ⟩, thus

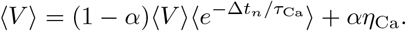

Solve for ⟨*V* ⟩:

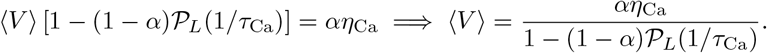

The time-average mean *µ*_Ca_ is obtained by averaging 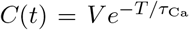 over the forward recurrence time *T* (time since last spike). The density of *T* is 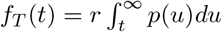, and

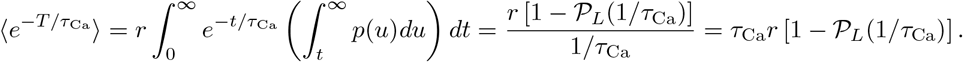

Thus,

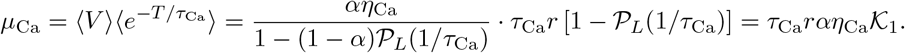

#### Second Moment of Calcium Concentration

Square the recurrence for *V*_*n*_:

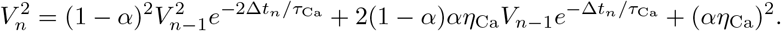

Take expectations in steady state for renewal processes, we have 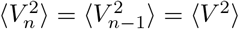, thus

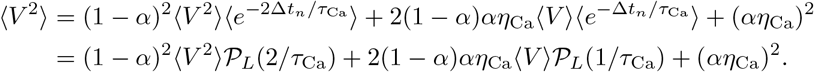

Substitute 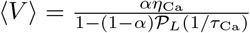 and rearrange:

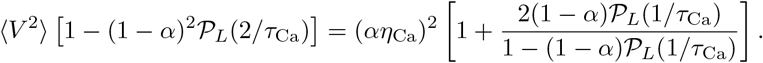

Simplify the right-hand side:

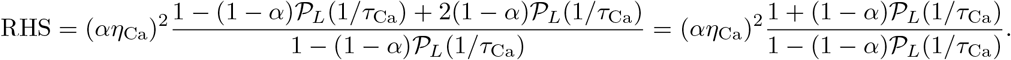

Thus,

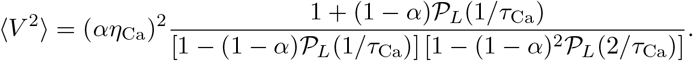

The second moment at arbitrary time *t* is 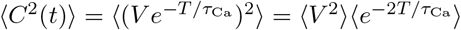 (by independence of *V* and *T*). The expectation 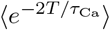 is:

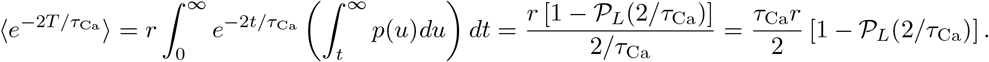

Therefore,

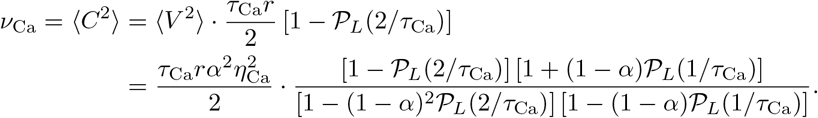

□

#### Poisson Limit Verification

For a Poisson process, 𝒫_*L*_(*s*) = *r/*(*r* + *s*). First, verify 𝒦_1_:

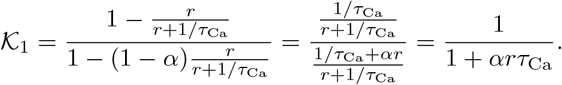

Thus 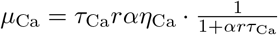, matching known Poisson result. For 𝒦_2_, we have

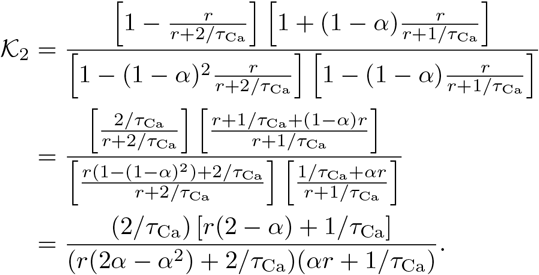

For *α* = 1 (no buffering), this simplifies to:

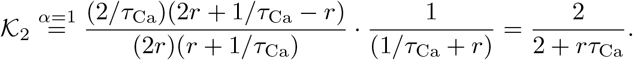

The full second moment is:

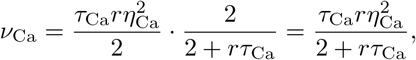

which matches the known result for *α* = 1 in the Poisson limit.

### 4.3 Divergent Encoding via Calcium Statistics

#### Theorem 2

(ISI-Variability Effects on Calcium Mean and Variance). *Let S*(*t*) *be a stationary renewal spike train with ISI family* {*p*(·; *θ*): *θ* ∈ Θ}, *each of mean* 1*/r, ordered by convex order:*

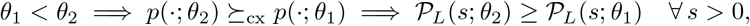

*where* 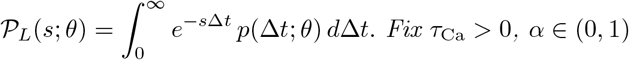, *and define*

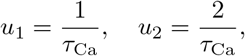

*and the dimensionless kernels*

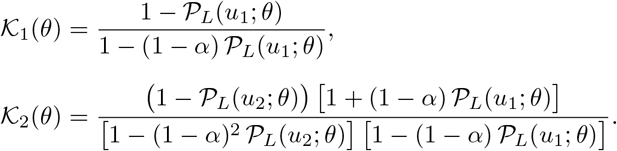

*Define the steady-state calcium moments*

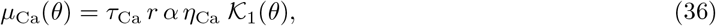

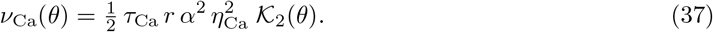

*Assume in addition the following* sufficient balanced buffering condition:

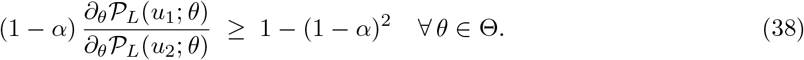

*Then*

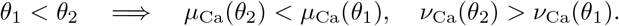

*Proof*.

1. **Laplace-transform ordering**. Convex order gives 𝒫_*L*_(*s*; *θ*_1_) ≤ 𝒫_*L*_(*s*; *θ*_2_) for all *s >* 0, hence *z*_*i*_(*θ*) = 𝒫_*L*_(*u*_*i*_; *θ*) is strictly increasing in *θ* for *i* = 1, 2.
2. **Monotonicity of** K_1_. Writing *z* = *z*_1_(*θ*),

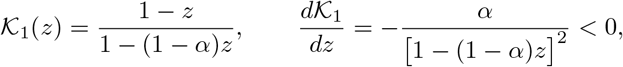

so 𝒦_1_(*θ*) strictly decreases.
3. **Monotonicity of** 𝒦_2_ **under the (sufficient) balancing condition in Eq. 38**. Let *z*_1_ = *z*_1_(*θ*), *z*_2_ = *z*_2_(*θ*). One computes

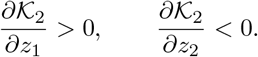

Therefore

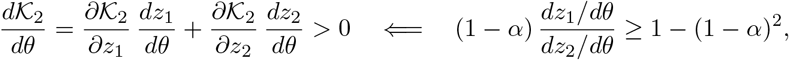

which is our sufficient balanced buffering condition. Hence 𝒦_2_(*θ*) strictly increases.
4. **Transfer to** *µ*_Ca_ **and** *ν*_Ca_. The prefactors in (36)–(37) are positive and independent of *θ*, so the monotonicity of 𝒦_1_ and 𝒦_2_ transfers directly to *µ*_Ca_(*θ*) and *ν*_Ca_(*θ*). □

### 4.4 Local Stability of the Calcium-based Homeostatic E-I Fixed Point

#### Theorem 3

(Fixed-Point Stability). *Let* 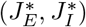 *be a fixed point of the closed-loop dynamics*

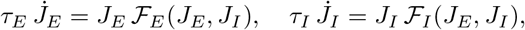

*corresponding equilibrium mean and variance of input as*

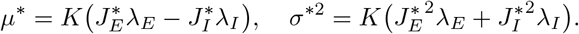

*Suppose at equilibrium*

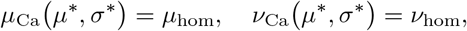

*and that the neuronal response functions r*(*µ, σ*) *and C*_*v*_(*µ, σ*) *satisfy*

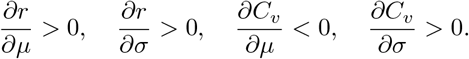

*Then the fixed point* 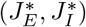 locally asymptotically stable.

*Proof*. Write

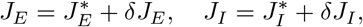

and linearize to first order:

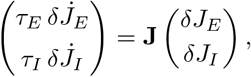

where the Jacobian **J** at 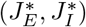 has entries

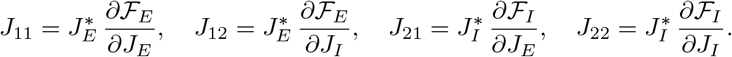

At equilibrium ℱ_*E*_ = ℱ_*I*_ = 0, and one finds

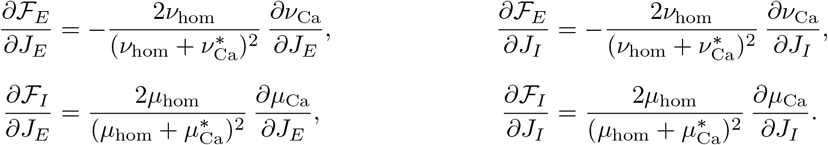

By the chain rule,

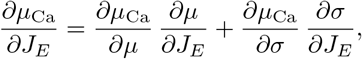

and similarly for the other partials, together with

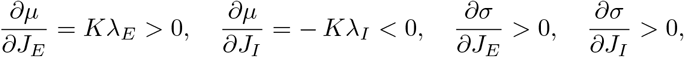

and under balanced buffering conditions stated in Theorem 2, we have *∂*_*µ*_*µ*_Ca_ *>* 0, *∂*_*σ*_*µ*_Ca_ ≥ 0, *∂*_*µ*_*ν*_Ca_ ≥ 0, *∂*_*σ*_*ν*_Ca_ *>* 0. Hence

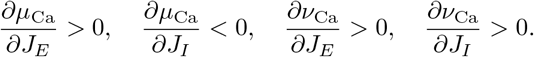

It follows that

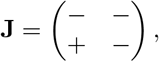

so tr **J** *<* 0 and det **J** *>* 0, implying both eigenvalues have negative real part. Therefore, the fixed point is locally asymptotically stable. □

### 4.5 Integrate-and-Fire Neuron Dynamics and its output spiking statistics

#### 4.5.1 Perfect Integrate-and-Fire (PIF)

Consider voltage dynamics *v* of a perfect Integrate-and-Fire (PIF) neuron [GKNP14] with membrane potential with input as in Eq. 1:

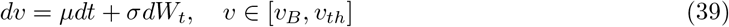

with reset to *v*_*r*_ upon reaching threshold *v*_*th*_ and *v*_*B*_ is reflective boundary condition. Using the Fokker-Planck equation for Eq. 39, the Laplace transform of ISI density is given by:

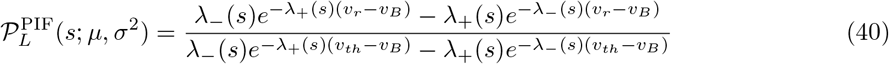

where 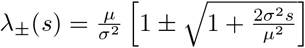. Thus, the output firing rate is

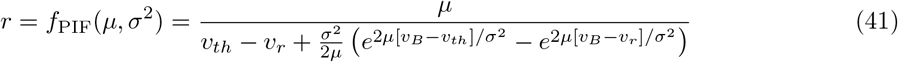

and the ISI coefficient of variation is given by

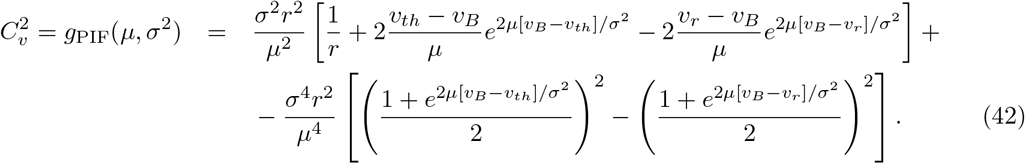

#### 4.5.2 Leaky Integrate-and-Fire (LIF)

Consider voltage dynamics *v* of a Leaky Integrate-and-Fire (LIF) neuron with membrane time constant *τ*_*m*_ and resting potential *v*_*l*_, driven by input as in Eq. 1:

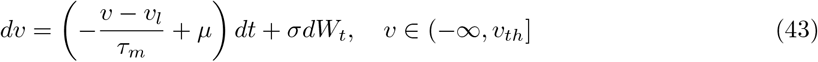

with reset to *v*_*r*_ upon reaching threshold *v*_*th*_. The stationary mean voltage is *v*_*∞*_ = *v*_*l*_ + *µτ*_*m*_. Using the Fokker-Planck equation for Eq. 43, the Laplace transform of the ISI density is given by [Bru00]:

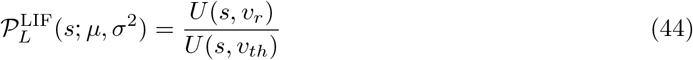

where *U* (*s, v*) is defined via parabolic cylinder functions [BH99]:

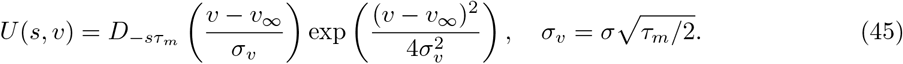

Here, *D*_*ν*_ (*z*) is the parabolic cylinder function. The output firing rate is [Tuc88]:

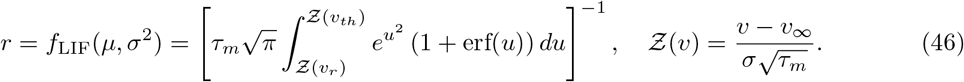

The ISI coefficient of variation *C*_*v*_ is given by [RS79]:

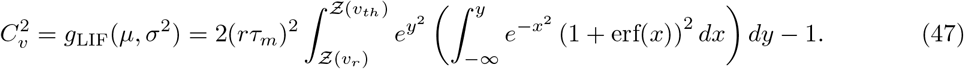

#### 4.5.3 Mean-Field Theory of Excitatory-Inhibitory Networks with External Input

We consider an E-I network where each neuron *i* in population *A* ∈ {*E, I*} receives inputs from populations *B* ∈ {*E, I, X*}, with *X* denoting an external Poisson drive at fixed rate *r*_*X*_. In the Erdős– Rényi connectivity model, the number of synapses from population *B* ∈ {*E, I, X*} onto neuron *i* in population *A* ∈ {*E, I*} is

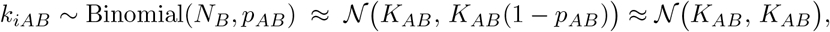

where *K*_*AB*_ = *N*_*B*_*p*_*AB*_ is the mean in-degree, and the Gaussian approximation holds for large *K*_*AB*_ and sparse connectivity (*p*_*AB*_ ≪ 1).

Under the diffusion approximation, synaptic inputs to neuron *i* in population *A* are characterized by mean *µ*_*iA*_ and variance 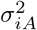:

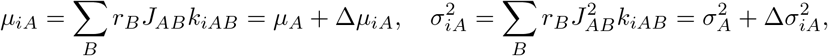

with population averages (*K* denotes a mean in-degree, assumed uniform for all populations *A* and *B*)

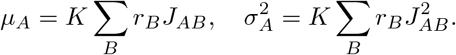

The quenched fluctuations Δ*µ*_*iA*_ and 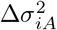 are zero-mean Gaussian random variables with variances

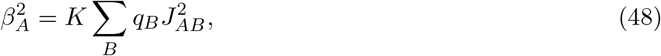

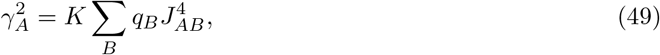

where 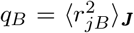 is the population and disorder averaged of the firing rates second moment for all *j* ∈ *B*.

##### Joint Distribution of Quenched Fluctuations

The covariance between Δ*µ*_*iA*_ and 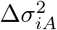 is:

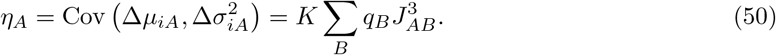

The joint distribution is bivariate Gaussian:

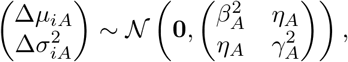

##### Self-Consistent Mean-Field Equations

Firing rates follow from the transfer function:

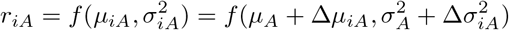

yielding population-averaged statistics:

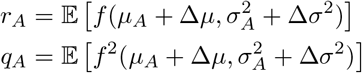

where expectation is over the bivariate Gaussian 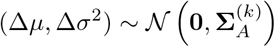 with covariance matrix

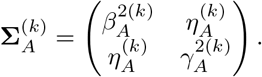

The system is solved iteratively:

1. Initialize 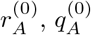
2. Compute statistics:

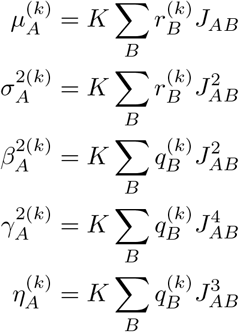
3. Update rates via Gaussian integration:

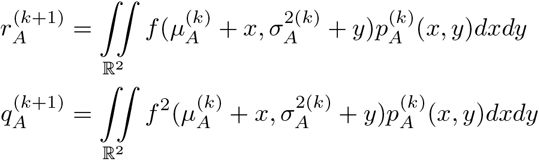

where 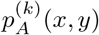 is the bivariate Gaussian density:

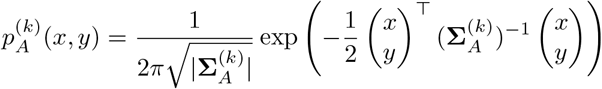
4. Repeat until convergence (∥**r**^(*k*+1)^ − **r**^(*k*)^∥ *< ϵ*)

##### Emergence of the balanced state

In systems where 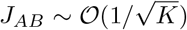, *η*_*A*_ and *γ*_*A*_ are of the lower order and can be ignored. We will show below that if the structure of *J*_*AB*_ allows *µ*_*A*_ also become 𝒪 (1), and in such a system, the convergence of all statistics is guaranteed, provided the stability of the system [vVS96]. This state is called the balanced state following the seminal work in [vVS96, vVS98]. In the balanced state, system-averaged temporal correlations are also weak in a wide range of network topologies, thus diffusion approximation of the input is admissible [Far25]. Furthermore, the existence and the stability of the balanced state indicate that the firing rates and spatial-temporal variances in the system remain non-zero and finite.

#### 4.5.4 Thermodynamic Limit and Self-Consistent Solution for the Balanced State

Following van Vreeswijk & Sompolinsky in [vVS98, vVS05], in the balanced regime, synaptic weights scale as

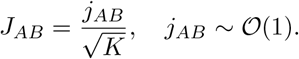

The total synaptic input to a neuron in population *A* decomposes into a power series [vVS05] as

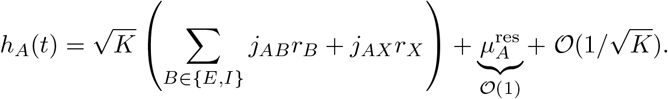

Balance van Vreeswijk condition requires cancellation of the 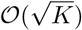 terms in thermodynamic limit:

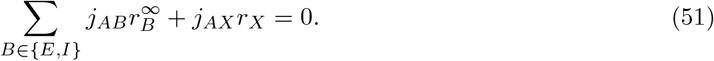

The existence of the balance state requires that the firing rates at the thermodynamic limit 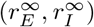 then satisfy:

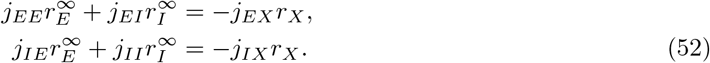

The balance-determined rates 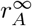 are first obtained analytically from Eq. (52). As 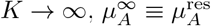 which must be determined self-consistently. Note that the stability of the balance state requires an inhibitory population feedback to stabilize the system faster than excitation dynamics [vVS98]. Provided the stability, we determine the self-consistent solution of the system at the balanced state.

##### Self-Consistent Residual Mean

The quenched fluctuation 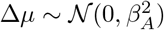 has statistics:

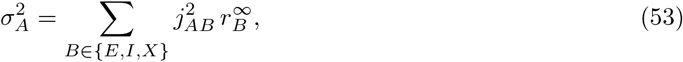

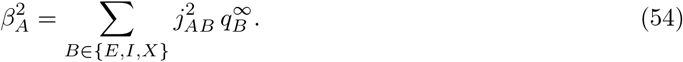

The self-consistent firing rate satisfies:

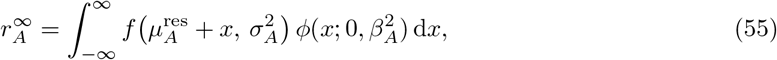

where 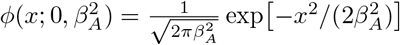. The second moment is

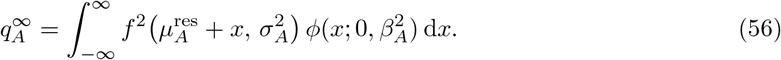

##### Numerical Solution Algorithm

1. **Solve for rates:** Obtain 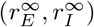 from Eq. (52).
2. **Initialize:**

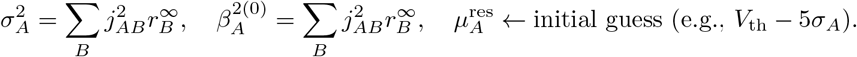
3. **Iterate until convergence:**

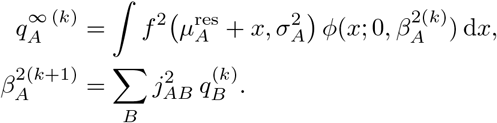
4. **Adjust residual mean:** Solve for 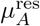 via root-finding (e.g. bisection) so that

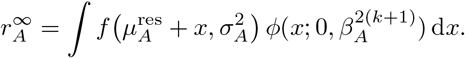
5. **Check convergence:** Terminate when 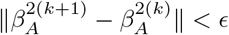 and 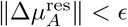.

## Acknowledgments

This work is dedicated to the memory of Carl van Vreeswijk, whose visionary insights into balanced network theory laid the foundation for much of modern theoretical neuroscience. Carl conceived the idea of the multidimensional calcium signal and performed the initial analytical calculations for the perfect integrate-and-fire (PIF) neuron model in 2021. His mentorship, curiosity, and profound intellectual generosity continue to inspire this work and the broader community he helped shape.

FF thanks Alex Reyes, David Hansel, and Rainer Engelken for fruitful discussions and comments on the manuscript. FF’s work was supported by the Deutsche Forschungsgemeinschaft (Grant No. FA1316/5-1).

